# Non-duality in brain and experience of advanced meditators – Key role for Intrinsic Neural Timescales

**DOI:** 10.1101/2025.04.21.649758

**Authors:** Saketh Malipeddi, Arun Sasidharan, Bianca Ventura, Rahul Venugopal, Clemens Christian Bauer, Prejaas K.B. Tewarie, P.N. Ravindra, Seema Mehrotra, John P John, Balachundhar Subramaniam, Steven Laureys, Bindu M Kutty, Georg Northoff

**Affiliations:** Centre for Consciousness Studies, Department of Neurophysiology, NIMHANS, Bengaluru, Karnataka, India; The Royal’s Institute of Mental Health Research & University of Ottawa, Brain and Mind Research Institute, Centre for Neural Dynamics, Faculty of Medicine, University of Ottawa, Canada; Northeastern University & McGovern Institute for Brain Research, MIT, USA; Joint International Research Unit on Consciousness, CERVO Brain Research Centre, Laval University, Canada; Sir Peter Mansfield Imaging Center, School of Physics, University of Nottingham, United Kingdom; Clinical Neurophysiology Group, University of Twente, Netherlands; Department of Clinical Psychology, NIMHANS, Bengaluru, Karnataka, India; Multi-modal Brain Image Analysis Laboratory, Department of Psychiatry, NIMHANS, Bengaluru, Karnataka, India; Sadhguru Center for a Conscious Planet, Beth Israel Deaconess Medical Center, Harvard Medical School, Boston, MA, United States; Joint International Research Unit on Consciousness, CERVO Brain Research Centre, Laval University, Canada; GIGA Consciousness Research Unit and Coma Science Group, Liège University, Belgium; International Consciousness Science Institute, Hangzhou Normal University, Hangzhou, China

**Author notes:** Equal contribution (Bindu M Kutty, Georg Northoff).

## Abstract

Distinguishing between self (internal) and environment (external) is fundamental to human experience, with ordinary waking consciousness structured around this duality. However, contemplative traditions describe non-dual states where this distinction dissolves. Despite its significance, the neural basis of non-duality remains underexplored. Using psychological questionnaires for non-duality experience and EEG-based intrinsic neural timescales as measured by the autocorrelation window (ACW), we studied non-duality in advanced meditators, novice meditators, and controls. All subjects underwent breath-watching meditation (internal attention) and a visual oddball cognitive task (external attention); this allowed us to conceptualize non-duality as a lack of distinction between internal and external attention. Our key findings include: (a) advanced meditators report greater experience of non-duality during breath-watching (psychological scales), (b) EEG-based ACW is longer during internal attention (breath watch) than external attention (oddball task) in all subjects taken together, (c) advanced meditators show no such distinction with equal duration of their ACW during both internal and external attention (we replicated this finding in another dataset of expert meditators); (d) the advanced meditators’ internal-external ACW difference correlated with their experience of the degree of non-duality (psychological scales) during internal attention. Together, these findings suggest that the brain’s intrinsic neural timescales during internal and external attention play a key role in mediating the experience of non-duality in advanced meditators.

## Introduction

Duality is a fundamental aspect of human experience, characterized by the distinction between internal and external phenomena. The internal phenomena are associated with self and self-related processing, while the external realm pertains to the world ^1–4^. Ordinary states of consciousness are structured around this internal-external distinction, amounting to a dual organization of their mental life.

This is different in non-ordinary states of consciousness, like meditation. These show that human consciousness is not confined to this dualistic framework; it also includes states that transcend such divisions as manifest in non-dual awareness ^5–14^. Deeply rooted in many contemplative, philosophical and spiritual traditions as diverse as Advaita Vedanta, Kashmiri Shaivism, Mahayana, and Vajrayana Buddhist schools ^8,12,15–17^, the experience of non-dual awareness leads to the dissolution of the internal and external duality ^6,12,14,18–24^. A defining feature of such non-dual states is “non-representational reflexivity”, which is the capacity to be aware without engaging in conceptual representation ^6,9^. In other words, it has been described as a state of pure awareness that exists beneath all mental constructs, such as categories, schemas, or metacognition ^9,11,25^. An emerging body of literature shows that advanced meditators report experiencing such non-dual states ^5,6,10,11,18,20,21,24,26–29^.

If non-duality represents a fundamental aspect of human experience, it will likely have distinct neural signatures. Preliminary evidence suggests that the central precuneus, a part of the default mode network (DMN), and its interactions with prefrontal regions may play a key role in mediating non-dual awareness ^5,6^. Research also shows neural synchronization of internal and external processing during advanced meditation with topographic re-organization of activity in the default mode network (DMN) (related to internal processing ^3,30–32^) and the central executive network (CEN) (related to external processing ^33–35^) in relation to the experience of non-duality ^11^. Given these findings in the spatial domain of the brain and its networks, how non-dual experiences manifest in its temporal features remains to be explored.

Recent evidence suggests that the brain is organised along a unimodal-transmodal hierarchy of intrinsic neural timescales (INTs) ^36–43^. INTs allow for the processing of stimuli characterized by different temporal signatures, i.e., different sets of timescales. Further, research shows that these INTs provide temporal receptive windows (analogous to spatial receptive fields in the visual cortex ^44–47^) that segregate and integrate inputs of varying durations ^39,48–55^. These INTs are measured by the auto-correlation function (ACF), which is a function “*that correlates a signal with copies of itself that are temporally shifted with a series of lags* ^37,38,41,52^.” It is typical in research to report the autocorrelation window (ACW), defined as “*the length of time at the moment when the ACF decays to 50% of its maximum value (ACW-50)* ^52,56–58^.” INTs have been involved in both internal and external tasks ^1,36–38,57,59,60^. Internal tasks have been associated with longer durations or timescales ^37,38,57,60^, while external tasks exhibit shorter duration or timescales ^37,61^. INTs have been shown to play a role in perception, behavior, cognition, consciousness, and a sense of self ^36–38,41,56,57,59,60^. Further, INTs have been shown to be sensitive to distinct attentional processes. For instance, meditative practices with broader attentional engagement, such as Shoonya, exhibit longer timescales, whereas practices with narrower attentional engagement, such as mantra meditation, show shorter timescales ^62^. This leaves open the role of INTs in non-duality, which can be observed across different meditative practices ^5,6,10,11,18,20,21,24,26–29,63^. Addressing this is the primary goal of our study.

In this study, we conceptualize duality in terms of internal and external processing, which we operationalized through distinct attention tasks: breath-watching meditation (internal attention) and a visual oddball cognitive task (external attention). Consequently, we operationalize non-duality as a reduction in the distinction between internal and external attention on both psychological (experience of non-duality) and neural (lack of difference in ACW between internal and external attention). To this end, we study how neural timescales in advanced meditators differ from those in novice meditators and controls during an internal-attention breath-watching task and an external-attention cognitive task (visual oddball); this serves the purpose to explore how neural changes in the internal-external tasks relate to the participants’ experience of non-duality at the psychological level. Besides the main data set, we also include a replication data (Supplement) where the main results are replicated.

Our study has three key aims: (1) characterizing non-duality at the psychological level using standard psychological scales, (2) identifying its neural markers and specifically its timescales by measuring EEG-based ACW, and (3) investigating the relationship between subjective experience (scales) and neural timescales (ACW) of non-duality.

Our first specific aim was to investigate non-duality at the psychological level among study participants. Based on previous literature ^5,6,10,11,18,20,21,24,26–29^, we hypothesized that advanced meditators would exhibit higher levels of the experienced state of non-duality and a fundamental tendency to not align themselves with either internal or external realms, e.g., non-attachment ^64–66^. Moreover, we hypothesized that non-attachment, as a trait feature, would correlate with state-level non-duality, reflecting the experienced state during meditation. Our second aim was to examine non-duality at the neural level in advanced meditators and controls during breath-watching (internal attention) and visual oddball tasks (external attention). We hypothesized that advanced meditators would be in a state of non-dual awareness and thus exhibit similar timescales during both the internally directed breath-watching meditation and the externally directed visual oddball task. Our third specific aim was to address whether these neural measures of non-duality, e.g., internal-external ACW difference, correlate with subjective measures of non-duality. We hypothesized that lesser differences in ACW between internal-attention breath-watching and external-attention cognitive task would relate to higher degrees in the experience of non-duality at the psychological level (Figure 1).

**Fig. 1:**
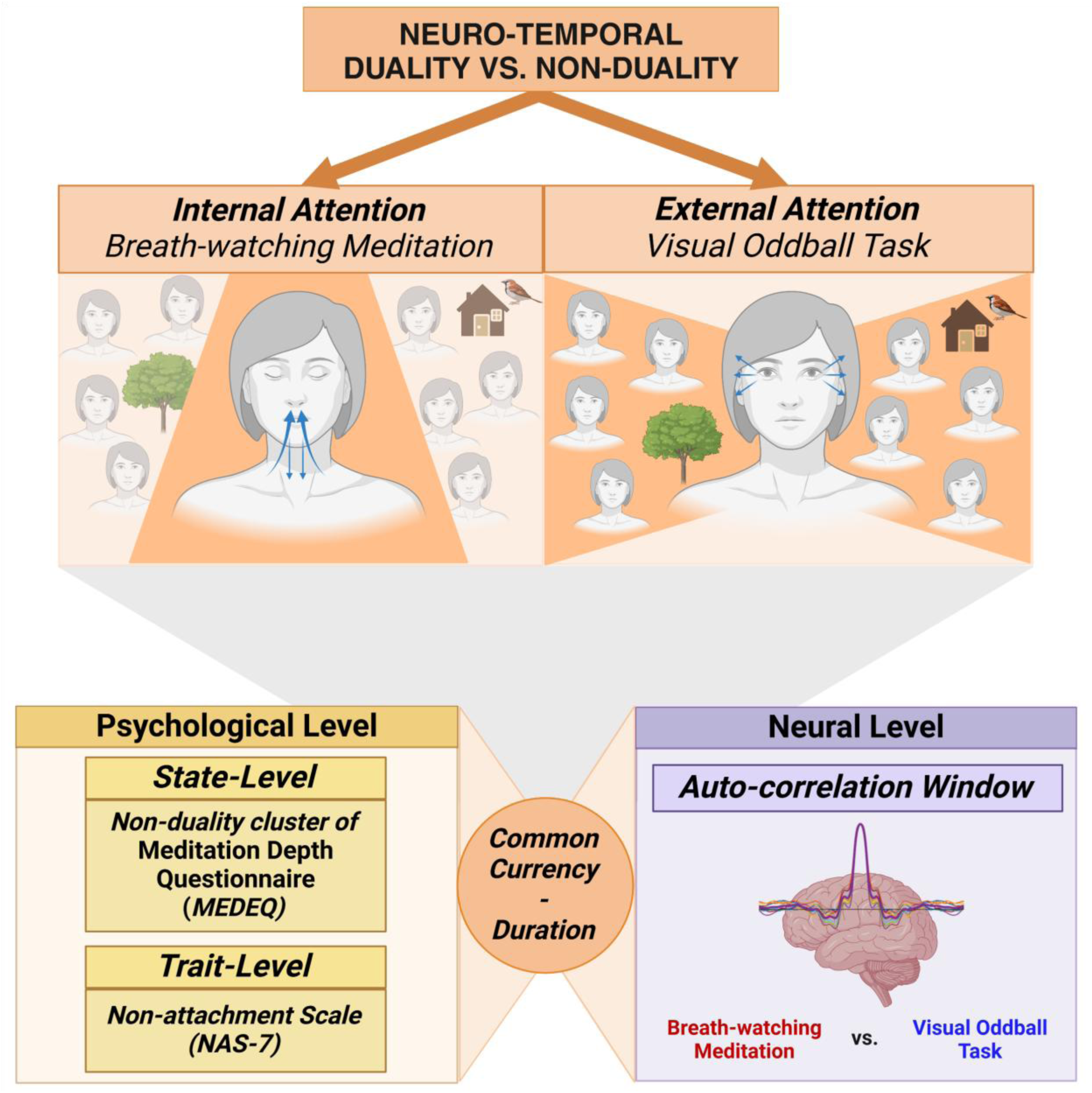
Hypothesis. The image illustrates how the study conceptualizes duality as the distinction between internal and external processing, operationalized through two attention tasks: breath-watching meditation (internal attention) and a visual oddball task (external attention). Non-duality is thus conceptualized as a reduction in this internal–external distinction. Duality and non-duality are examined at both the psychological level (using the non-duality cluster of the Meditation Depth Questionnaire ^67^ and the Non-Attachment Scale ^68^) and the neural level (using EEG-based autocorrelation window during internal and external attention tasks). Temporal duration serves as a common currency linking mind and brain as it is supposedly shared by both neural and mental states, e.g., brain and experience ^1^.

## Results

### Psychological features of non-duality

#### Meditation depth

To assess how well the study participants performed the breath-watching meditation, we investigated the differences in meditation depth, as measured by the meditation depth questionnaire, among our three groups. As illustrated in Figure 2(a), the results show a significant difference in meditation depth (χ^2^_Kruskal-Wallis_(2) = 25.42, *p* < 0.0001, ɛ^2^_ordinal_ = 0.37, CI_95%_[0.25, 1.00]) between the three groups. Advanced meditators exhibited the highest levels of meditation depth, followed by novices, with controls reporting the lowest levels. The effect sizes were large. Post-hoc comparisons showed significant differences between advanced meditators and controls (*p*_Holm-adj_ < 0.0001), between advanced and novice meditators (*p*_Holm-adj_ = 0.03), and between novices and controls (*p*_Holm-adj_ = 0.01). As expected, we found a significant positive correlation between lifetime hours of meditation and meditation depth (*Ρ_Spearman_* = 0.60, *p* < 0.0001) (Supplement Figure 1a).

**Fig. 2:**
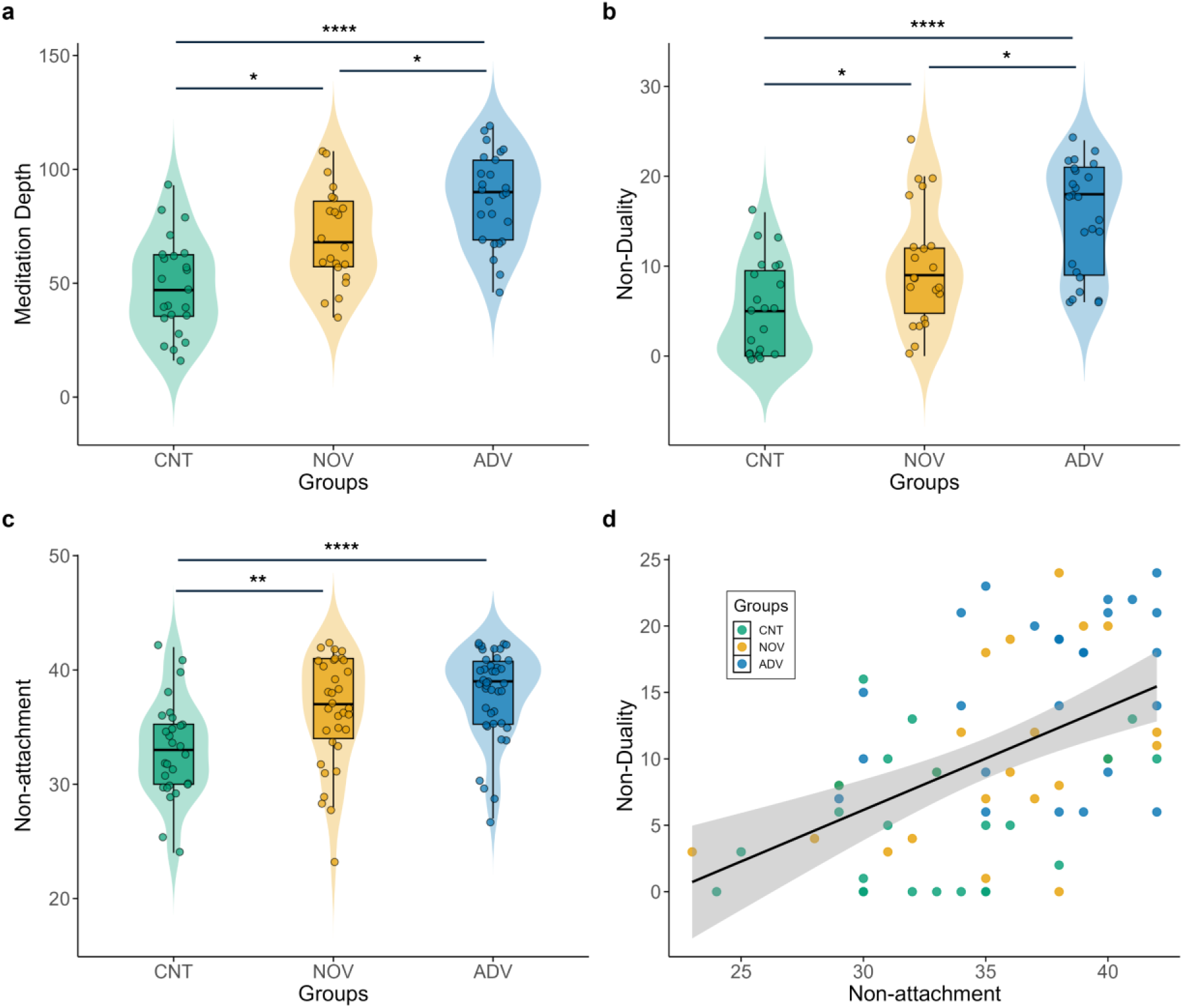
Group differences in meditation depth, non-duality and non-attachment. **(a)** Box-violin plots depicting group differences in Meditation Depth among control (CNT), novice (NOV), and advanced (ADV) meditators. Advanced meditators (blue) showed the highest Meditation Depth scores, followed by novice meditators (orange) and control participants (teal) (χ^2^_Kruskal-Wallis_(2) = 25.42, p < 0.0001, ɛ^2^_ordinal_ = 0.37, CI_95%_[0.25, 1.00], n = 70; CNT = 23, NOV = 22, ADV = 25). **(b)** Box-violin plots depicting group differences in non-duality, demonstrating increased non-dual experiences in advanced meditators (χ^2^_Kruskal-Wallis_(2) = 22.94, p < 0.0001, ɛ^2^_ordinal_ = 0.33, CI_95%_[0.21, 1.00], n = 70; CNT = 23, NOV = 22, ADV = 25). **(c)** Box-violin plots showing non-attachment differences across groups, with higher scores in meditators (χ^2^ (2) = 18.42, p < 0.0001, ɛ^2^ = 0.18, CI_95%_[0.10, 1.00], n = 103; CNT = 28, NOV = 33, ADV = 42). **(d)** Scatterplot illustrating the positive relationship between Non-Duality and Non-Attachment across groups (Ρ_Spearman_ = 0.51, p < 0.0001, n = 70). Significance levels: *p < 0.05, **p < 0.01, ****p < 0.0001.

Together, these findings support and validate that our advanced meditators were indeed advanced, based on their lifetime meditation hours and their meditation depth during breath-watching meditation. Correspondingly, lifetime hours and meditation depth correlated with each other. Hence, our advanced meditators are truly advanced in both objective (lifetime hours) and subjective (meditation depth) terms, distinguishing them from novices, who, in turn, were different from the controls. This further validates our measure of meditation depth with regarding the breath-watching meditation.

#### Non-duality in advanced meditators

Non-duality is a state of experience where subjects no longer differentiate between their internal reality (e.g., thoughts and the self) and external reality (e.g., events and objects in the environment). By measuring the state of non-duality during meditation, i.e., breath-watching, with the well-validated non-duality cluster of the meditation depth scale, we could compare our three groups in terms of their degrees of non-dual experience during the preceding meditation. We supposed the experience/perception of non-duality to be higher in the advanced meditators than in the other two groups. This hypothesis was confirmed: as illustrated in Figure 2(b), the results show a significant difference in the experience of non-duality during breath-watching (χ^2^_Kruskal-Wallis_(2) = 22.94, *p* < 0.0001, ɛ^2^_ordinal_ = 0.33, CI_95%_[0.21, 1.00]) between the three groups. Advanced meditators exhibited the highest levels of non-duality, followed by novices, with controls reporting the lowest levels. The effect sizes were large. Post-hoc comparisons for non-duality showed significant differences between advanced meditators and controls (*p*_Holm-adj_ < 0.0001), between advanced and novice meditators (*p*_Holm-adj_ = 0.04), and between novices and controls (*p*_Holm-adj_ = 0.04). Further, we found a positive correlation between non-duality and lifetime hours of meditation (*Ρ_Spearman_* = 0.57, *p* < 0.0001) (Supplement Figure 1b).

Our results show that advanced meditators experience high degrees of non-duality, significantly greater than those of novices and controls. Moreover, the experience of non-duality correlated with the lifetime hours, further supporting the validity of our non-duality measure. Accordingly, non-duality emerges as a key feature of the breath-watching state, particularly in advanced meditators and, to a lesser extent, in novices and controls.

#### Non-attachment and its relation to non-duality

We measured non-duality experience directly in relation to the meditative state during breath-watching, which is associated with lifetime meditation hours. This raises the question of whether the experience of non-duality is not only a state-dependent feature but also a trait feature in the advanced meditators. One key feature of the non-dual state is the individuals’ ability to refrain from identifying with, aligning with, or attaching to any form of content, including internal thoughts or feelings and external environmental stimuli. This ability can be measured with the non-attachment scale, which quantifies the subjects’ capacity to not attach or identify themselves with any content. Results (Figure 2(c)) show a significant difference between the groups in non-attachment (χ^2^_Kruskal-Wallis_(2) = 18.42, *p* < 0.0001, ɛ^2^_ordinal_ = 0.18, CI_95%_[0.10, 1.00]). The effect sizes were large. Advanced meditators reported the highest scores for non-attachment. Post-hoc comparisons for non-attachment showed significant differences between advanced meditators and controls (*p*_Holm-adj_ < 0.0001) and between novices and controls (*p*_Holm-adj_ < 0.01). We also found a significant positive correlation between non-attachment and lifetime hours of meditation (*Ρ_Spearman_* = 0.44, *p* < 0.0001) (Supplement Figure 1c). Finally, as expected, we found a significant positive correlation between non-attachment and non-duality (*Ρ_Spearman_* = 0.51, *p* < 0.0001) (Figure 2(d)), indicating that higher levels of non-attachment were associated with greater degrees of non-duality. Accordingly, our results show that the subjects’ capacity for non-attachment relates to their non-duality during meditation.

Together, our findings demonstrate that advanced meditators also exhibit greater trait capacity for non-attachment to both internal and external mental content. Furthermore, the strong correlation between non-attachment and non-duality supports the idea that non-attachment is a key characteristic of non-duality, with higher levels of non-attachment (as a trait) associated with greater degrees of non-duality (as a state). Hence, our data strongly support the central role of non-duality and non-attachment as a core feature of the breath-watching experience in advanced meditators, distinguishing them from both novices and controls.

### Neuronal features

#### Neuro-temporal duality - Longer auto-correlation window (ACW) during internal (breath watch) than external (visual oddball) attention across all subjects

Non-duality refers to the absence of a distinction between the internal self and the external environment, thereby overcoming their duality in our common experience ^6,8^. This concept can be operationalized at the neural processing level by comparing the difference in EEG-based ACW duration between an internal-attention breath-watching meditation and an external-attention cognitive task. We hypothesized that the difference in ACW duration between internal and external attention tasks would indicate the level of dual (*‘neuro-temporal duality’*) or non-dual (*‘neuro-temporal non-duality’*) processing at the neural level. To test this, we first combined all the groups. We examined the differences in ACW mean between breath-watching and cognitive task conditions using an unpaired t-test, thus assessing the overall neural difference between internal (breath-watching meditation) and external (visual oddball task) processing (Figure 3(a)). Our results show a statistically significant difference, with a very large effect size, between the internal-oriented breath-watching and the external-oriented cognitive task (*t*(163.52) = 6.09, *p* < 0.0001, *g_Hedges_* = 0.91, CI_95%_[0.60, 1.21]). In other words, the ACW mean becomes shorter and thus narrower (in its temporal cross-correlation between distant time points) during an externally oriented condition (visual oddball cognitive task). In contrast, it is longer and thus wider during an internally-oriented condition (breath-watching meditation).

**Fig. 3:**
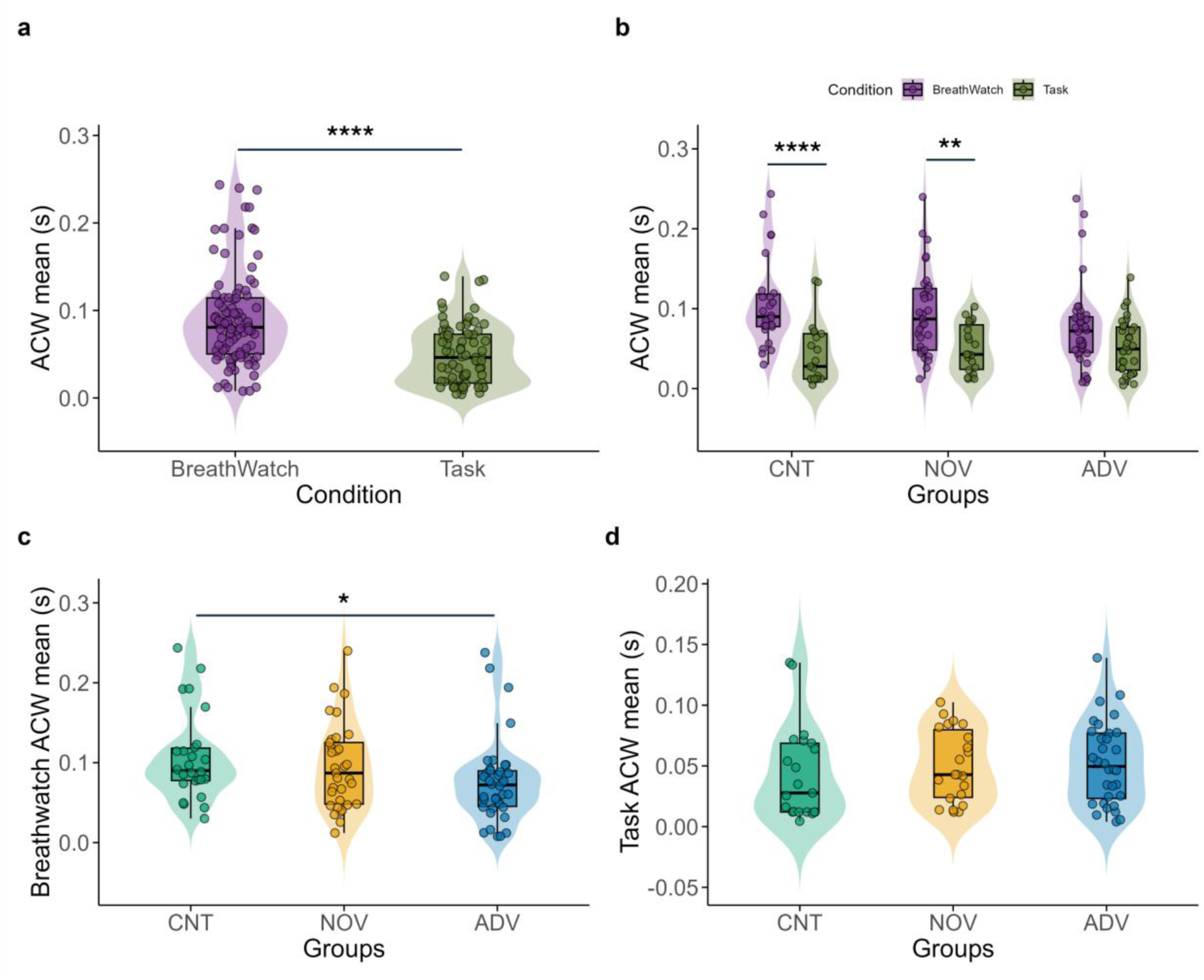
Autocorrelation window (ACW) across conditions and groups. **(a)** Difference in ACW mean between internal-attention breath-watching meditation and external-attention cognitive task (all groups combined). ACW becomes shorter and narrower during the external task while it becomes longer and wider during the internal-attention breath-watching (t(163.52) = 6.09, p < 0.0001, g_Hedges_ = 0.91, CI_95%_[0.60, 1.21], n = 172; BreathWatch = 97, Task = 75)). **(b)** Group-wise ACW mean for both conditions, where CNT and NOV groups exhibit significantly higher ACW mean in the BreathWatch condition compared to the Task condition (CNT: t(70) = 4.804, p = 0.0001; NOV: t(70) = 3.782, p = 0.0042), whereas ADV do not show any significant difference between these two conditions (t(70) = 1.770, p = 0.4918). **(c)** Comparison of ACW mean during BreathWatch across groups (CNT, NOV, ADV), showing a significantly lower ACW in the advanced meditators (χ^2^_Kruskal-Wallis_(2) = 6.59, p = 0.04, ɛ^2^_ordinal_ = 0.07, CI_95%_[0.00981, 1.00], n = 97; CNT = 26, NOV = 33, ADV = 38)). **(d)** No significant differences in ACW mean between groups during the Task condition (χ^2^_Kruskal-Wallis_(2) = 2.23, p = 0.33, ɛ^2^_ordinal_ = 0.03, CI_95%_[0.00475, 1.00], n = 75; CNT = 21, NOV = 22, ADV = 32). Significance levels: *p < 0.05, **p < 0.01, ****p < 0.0001. CNT: Controls, NOV: Novice meditators, ADV: Advanced meditators.

Together, these results confirm our hypothesis that the duality of the internal (breath-watching) and the external (cognitive task) on the psychological level is also manifest on the neural level in terms of different processing durations, i.e., timescales, as indicated by longer ACW during breath-watching and shorter ACW during the cognitive task. This further suggests that ACW during breath-watching and the cognitive task can serve as a neural proxy for duality vs. non-duality. Consequently, one may thus want to speak of ‘*neuro-temporal duality*’ featuring the different processing durations – i.e., timescales - of internally (breath-watching) and externally (visual oddball) directed attention.

#### Neuro-temporal non-duality I - No difference between internal and external ACW in advanced meditators

Next, we assessed how meditation experience influences this ACW-based neuro-temporal duality in processing external and internal attention conditions. To this end, a mixed ANOVA was conducted to examine the effects of meditation experience (group: advanced, novice, control) and condition (breath-watching, visual oddball) on the dependent variable, ACW means (Table 1). The analysis revealed a significant main effect of condition, *F*(1, 70) = 37.84, *p* < 0.0001, indicating that the ACW mean differed significantly between the breath-watching and cognitive task conditions. However, the main effect of the group was not significant, *F*(2, 70) = 0.74, *p* = 0.48, suggesting that lifetime meditation hours and exposure alone are unlikely to account for differences in ACW. In contrast, there was a significant interaction effect between group and condition, *F*(2, 70) = 3.76, *p* = 0.028, indicating that the effect of condition varied across groups. This finding emphasizes the relationship between ACW and the actual state (i.e., breath-watching vs. visual oddball cognitive task) rather than a purely trait-based effect of meditation experience (i.e., a group independent of condition).

**Table 1:** Breath-watching meditation vs Cognitive Task - Results of the Mixed ANOVA Examining the Effects of Group and Condition on ACW Mean.

| Effect | Sum Sq | df_num | Error SS | df_den | F value | p-value |
| --- | --- | --- | --- | --- | --- | --- |
| Intercept | 0.68911 | 1 | 0.18898 | 70 | 255.249 | $p < 2.2e-16$ **** |
| Groups | 0.004 | 2 | 0.18898 | 70 | 0.7411 | $p = 0.4803$ <sup>ns</sup> |
| Condition | 0.06274 | 1 | 0.11607 | 70 | 37.8391 | $p = 4.2e-08$ **** |
| Interaction (Groups<br>× Condition) | 0.01247 | 2 | 0.11607 | 70 | 3.7607 | $p = 0.0281$ * |

*This table presents the results of a mixed ANOVA assessing the main effects of Group (Controls, Novice Meditators, Advanced Meditators) and Condition (Breath-Watching Meditation, Cognitive Task) and their interaction on ACW mean. The table includes the sum of squares (Sum\_Sq), numerator degrees of freedom (df\_num), error sum of squares (Error\_SS), denominator degrees of freedom (df\_den), F-values (F\_value), and corresponding p-values (p\_value). Significance levels are denoted as: ns = Not significant ( $p \geq 0.05$ ), \* = $p < 0.05$ , \*\* = $p < 0.01$ , \*\*\* = $p < 0.001$ , \*\*\*\* = $p < 0.0001$ .*

Post-hoc pairwise comparisons were conducted to assess differences in ACW mean between breath-watching meditation and the visual oddball cognitive task within each group of meditators (advanced, novice, control) (Figure 3(b)). This difference was computed to test for the degree of duality (large ACW differences between breath-watching and visual oddball cognitive task) or non-duality (small ACW differences between breath-watching and visual oddball cognitive task) on the neuronal level. In advanced meditators, no significant difference in ACW mean was observed between breath-watching and the visual oddball cognitive task (t(70) = 1.770, *p* = 0.4918), indicating that their neural processing duration (i.e., ACW mean) remained similar across internal (breath-watching) and external (visual oddball cognitive task) domains. In contrast, novice meditators showed a significant ACW difference between internal and external tasks (t(70) = 3.782, *p* = 0.0042), suggesting distinct neural processing durations for internal and external tasks. Similarly, the control group exhibited a highly significant difference (t(70) = 4.804, *p* = 0.0001) (Figure 3(b)), suggesting a strong condition effect in this group. All p-values were adjusted using the Tukey method for multiple comparisons. Together, these results demonstrate a more non-dual neural expression of ACW in advanced meditators (i.e., no significant difference between breath-watching and the cognitive task). In contrast, novices and controls exhibited a more dual processing pattern (i.e., a significant ACW difference between conditions).

Next, given that the ACW is known to reflect temporal integration (longer ACW) and segregation (shorter ACW) ^38,60^, as entailed by both breath-watching and the visual oddball cognitive task (see the introduction for details on that), we examined group differences in ACW values for each condition separately. We observed that the ACW during breath-watching was significantly shorter in the advanced meditators than in both controls and novices (χ^2^_Kruskal-Wallis_(2) = 6.59, *p* = 0.04, ɛ^2^_ordinal_ = 0.07, CI_95%_[0.00981, 1.00]) (Figure 3(c)). In contrast, there was no significant difference in the ACW of the externally-oriented visual oddball cognitive task between the three groups (χ^2^_Kruskal-Wallis_(2) = 2.23, *p* = 0.33, ɛ^2^_ordinal_ = 0.03, CI_95%_[0.00475, 1.00]) (Figure 3(d)). In the behavioural measures of the visual oddball task, response times did not significantly differ across the three groups (Supplementary Figure 2a). However, response accuracy showed significant group differences (Supplementary Figure 2b), with advanced meditators demonstrating the highest accuracy compared to the other groups.

Together, we show that advanced meditators exhibit significantly less or lower ACW difference between internal (breath-watching) and external (oddball) conditions compared to both controls and novices. This effect is primarily driven by a shortening of the ACW during the internal condition (breath-watching). At the same time, it exhibits the same duration among all three groups during the external task. Accordingly, unlike the other two groups, the ACW in advanced meditators approximates and becomes similar to the ACW during the external condition. This indicates that both internal and external contents are processed similarly at the neural level, that is, they are processed with similar duration in their timescales. Therefore, ACW duration in advanced meditators no longer differs between internal and external processing, suggesting a higher degree of non-dual processing at the neural level -what we refer to as ‘*neuro-temporal non-duality*.’ Conversely, controls and novices exhibit greater differentiation in ACW duration between internal and external tasks, indicating more dual processing in the temporal dynamics of neural activity, thus reflecting a higher degree of neuro-temporal duality.

#### Neuro-temporal non-duality II – Is it related to the brain’s spontaneous activity?

We so far demonstrated that differences in the psychological state - dual vs. non-dual - relate to corresponding neural features in the brain’s processing duration, as measured by the ACW. Controls and novices exhibited different lengths in their ACW during internal and external tasks, reflecting a stronger degree of neuro-temporal duality. Conversely, advanced meditators no longer showed such ACW difference, reflecting a tendency towards neuro-temporal non-duality. This raises the question of where and how neuro-temporal differences in advanced meditators arise during internal (breath-watching) task.

To explore this, we investigated spontaneous neural activity during the resting state, which has been shown in previous papers to predict task-related differences in ACW ^38,61^. We first compared the ACW in the resting state among the three groups, finding no significant differences. Next, we compared the differences between the resting state and the internal task (breath-watching) across the three groups. A mixed ANOVA was conducted to examine the effects of meditation experience (Group: advanced, novice, control) and condition (breath-watching, resting state) on the dependent variable, ACW mean (Table 2) (Figure 4(a)). The analysis revealed a significant main effect of condition, *F*(1, 76) = 37.8391, *p* < 0.001, indicating that the ACW mean differed significantly between the breath-watching and resting state conditions. However, the group’s main effect was insignificant, *F*(2, 76) = 0.7411, *p* = 0.3760. Conversely, there was a significant interaction between group and condition, *F*(2, 76) = 3.76, *p* = 0.0298, indicating that the effect of condition varied across groups.

**Fig. 4:**
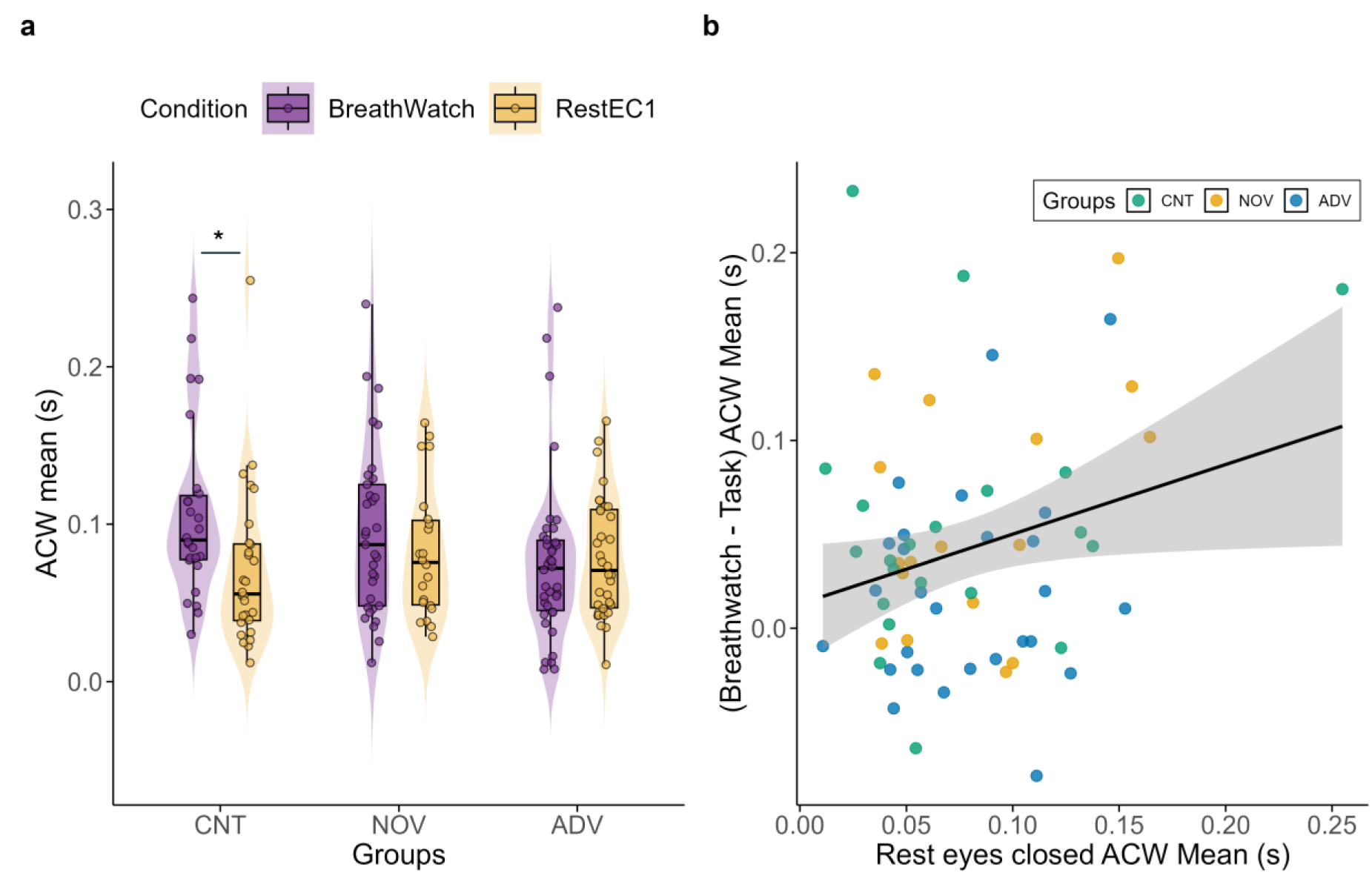
Autocorrelation window (ACW) mean duration for breath-watch vs rest. **(a)** The difference in ACW mean between internal-attention breath-watching meditation and eyes closed resting state. There is no significant difference between breath-watching and rest in novice (t(76) = 2.639, p = 0.1003) and advanced (t(76) = 0.053, p = 1.0000) meditators. In contrast, controls show significant differences (t(76) = 3.477, p = 0.0105) (**b)** Scatter plot illustrating the significant positive correlation between Rest ACW mean and (Breathwatch – Task) ACW mean (r = 0.26, p = 0.04, n = 65). Shaded areas represent 95% confidence intervals. CNT = Controls; NOV = Novice meditators; ADV = Advanced meditators.

**Table 2:** Breath-watching meditation vs Resting state - Results of the Mixed ANOVA Examining the Effects of Group and Condition on ACW Mean.

| Effect (Breath-watch vs Rest) | Sum_Sq | df_num | Error_SS | df_den | F_value | p_value |
| --- | --- | --- | --- | --- | --- | --- |
| Intercept | 1.21358 | 1 | 0.274508 | 76 | 255.249 | < 2.2e-16 **** |
| Groups | 0.00716 | 2 | 0.274508 | 76 | 0.7411 | 0.3760335 <sup>ns</sup> |
| Condition | 0.01742 | 1 | 0.097757 | 76 | 37.8391 | 0.0004341 *** |
| Interaction (Groups × Condition) | 0.00946 | 2 | 0.097757 | 76 | 3.7607 | 0.0298918 * |

*This table presents the results of a mixed ANOVA assessing the main effects of Group (Controls, Novice Meditators, Advanced Meditators) and Condition (Breath-Watching Meditation, Resting state) and their interaction on ACW mean. The table includes the sum of squares (Sum\_Sq), numerator degrees of freedom (df\_num), error sum of squares (Error\_SS), denominator degrees of freedom (df\_den), F-values (F\_value), and corresponding p-values (p\_value). Significance levels are denoted as: ns = Not significant ( $p \geq 0.05$ ), \* = $p < 0.05$ , \*\* = $p < 0.01$ , \*\*\* = $p < 0.001$ , \*\*\*\* = $p < 0.0001$ .*

Further, post-hoc pairwise comparisons were conducted to assess differences between breath-watching meditation and resting state within each group of meditators (Figure 4(a)). In the advanced meditators, no significant difference in ACW mean was observed between breath-watching and resting state (*t*(76) = 0.053, *p* = 1.0000). Similarly, in the novice meditators, no significant difference was found (*t*(76) = 2.639, p = 0.1003). However, in the control group, a significant difference was observed (*t*(76) = 3.477, p = 0.0105), suggesting a difference in ACW mean between breath-watching and resting state. Finally, we found a significant positive correlation of the resting state ACW with the internal (breath-watching)-external (visual oddball) task ACW differences (*r* = 0.26, *p* = 0.04) (Figure 4(b)). This suggests that the degree of duality (or non-duality) on the neuronal level (difference in ACW between internal and external) relates to the duration of the ACW in the resting state: longer duration in the latter is associated with larger differences in the former, entailing a higher degree of dual internal-external processing. In contrast, shorter ACW in the resting state leads to lower internal-external ACW differences, reflecting a more non-dual internal-external processing.

We also conducted the same analysis for the resting state-external task (visual oddball) differences among the three groups, which revealed a significant main effect of condition but no significant effects of group or group × condition interactions (Supplementary Table 1, Figure 3). Post-hoc comparisons indicated a marginally significant difference between the resting state and the external task (oddball) for controls (*p* = 0.08) (Supplementary Figure 3). In contrast, no significant differences were observed for novices or advanced meditators.

Overall, these results suggest that advanced meditators do not show significant ACW differences across the division of intrinsic (resting state), external (visual oddball task), and internal (breath-watching meditation) states; that is, for internal breath-watching vs. external task (oddball), resting state vs. breath-watching, and resting state vs. external task (oddball). In the advanced meditators, ACW duration remains consistent across all three states, whereas this is not the case in the other two groups. Hence, these findings further support the idea of *neuro-temporal non-duality* as a core feature of neural processing in advanced meditators. In contrast, controls exhibit significant differences across all conditions, supporting that their neural processing conforms to *neuro-temporal duality*.

#### Neuro-temporal non-duality in replication dataset

In an independent dataset of 43 expert meditators (see Methods), we replicated the neuro-temporal non-duality findings observed in our primary sample. There were no significant differences in ACW duration between breath-watching meditation, rest, and the visual oddball task (F(2, 64.68) = 0.06, *p* = 0.95; Supplementary Figure 4). This further supports the validity of our results.

### Linking psychological duality/non-duality to neuro-temporal duality/non-duality

#### Correlation between ACW and non-duality

How is the similarity of the ACW on the neural level related to the experience of the duality/non-duality on the psychological level? Our final step links ACW duration during internal breath-watching and external cognitive tasks, to the psychological measures of non-duality. To explore this, we investigated the relationship between the internal-external ACW duration difference (BW-Task) and the psychological scores of non-duality. Our results showed a significant negative correlation between non-duality and ACW (BW-Task) (*Ρ_Spearman_* = −0.31, *p* = 0.0227) (Figure 5(a)), indicating that a smaller internal-external ACW difference at the neural level is associated with a higher degree of non-duality at the psychological level.

**Fig. 5:**
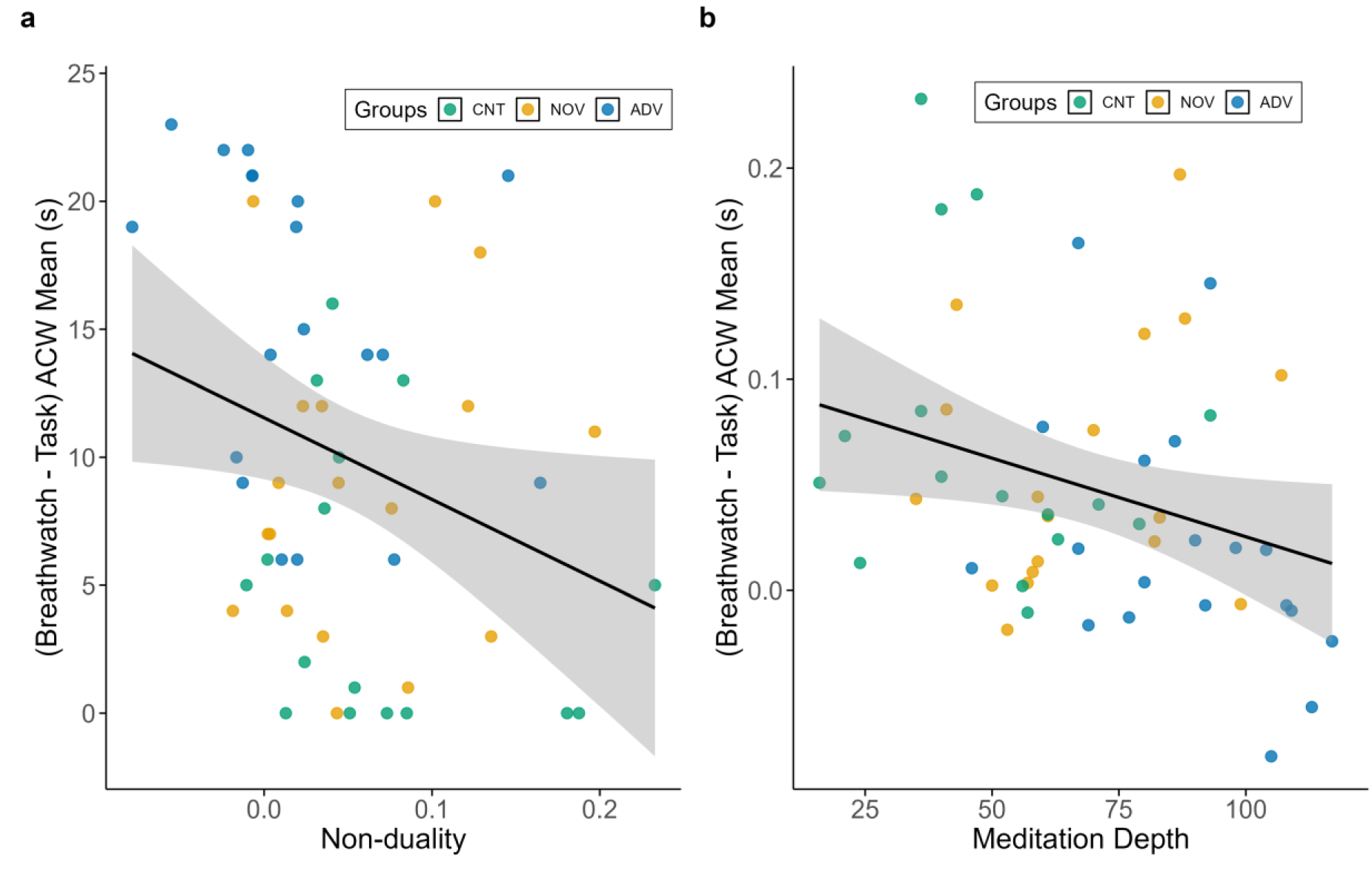
Correlation between ACW mean and non-duality. **(a)** Scatter plot illustrating the significant negative correlation between (Breathwatch – Task) ACW mean and non-duality (Ρ_Spearman_ = −0.31, p = 0.0227, n = 53). **(b)** Scatter plots depict the significant negative correlation between the ACW mean (Breath-watch – Task) and meditation depth (Ρ_Spearman_ = −0.29, p = 0.03, n = 53). Shaded areas represent 95% confidence intervals. CNT = Controls; NOV = Novice meditators; ADV = Advanced meditators.

These findings suggest that the neural measure of non-duality - specifically, the internal-external ACW duration difference - relates to non-duality on the psychological level. As mentioned in the previous section, the smallest difference (or lack thereof) in ACW between the dual conditions (breath-watching vs. visual oddball task) was observed in the advanced meditators, while the largest difference was observed in the control group, with novices falling in between.

In summary, we show that smaller ACW differences between the dual processing of the internal (breath-watching) and external (visual oddball task) conditions relate to higher non-duality scores on the psychological level. This supports a direct relationship between neuro-temporal duality/non-duality in the brain and the subjective experience of duality/non-duality on the psychological level.

Finally, we investigated whether the depth of meditation and lifetime meditation hours were associated with the difference in ACW mean between the internal attention (breath-watching) and external attention (visual oddball cognitive task) conditions. Consistent with our findings on non-duality, we observed a significant negative correlation of the difference-based ACW (Breath-watch – Task) with both meditation depth (*Ρ_Spearman_* = −0.29, *p* = 0.03) (Figure 5(b)) and lifetime hours of meditation (*Ρ_Spearman_* = −0.25, *p* = 0.03) (Supplementary Figure 2): a smaller internal-external ACW difference is associated with more meditation depth and more lifetime hours of meditation.

Taken together, our results demonstrate that the degree of similarity of internal-external tasks in their ACW duration in the three groups is related to key psychological measures, i.e., the experience of non-duality and meditation depth and lifetime meditation hours. These findings further support the central role of the brain’s neural processing duration, i.e., ACW, in the dual vs. non-dual processing of external and internal stimuli, thus modulating the psychological effects of meditation, including non-duality, meditation depth, and lifetime meditation hours.

## Discussion

Our research investigated the psychological and neural features of non-duality in advanced meditators, novice meditators, and controls. We studied non-duality using two tasks: breath-watching meditation for internal attention and a visual oddball task for external attention. Non-duality is defined as a reduction in the distinction between these two types of attention, examined both psychologically (the experience of non-duality) and neurally (similar ACW for both).

We discovered the following: a) advanced meditators experienced a greater sense of non-duality at the psychological level during breath-watching meditation compared to other groups; b) the duality between internal and external attention is also evident at the neural level: longer intrinsic neural timescales, measured by the autocorrelation window (ACW), were observed during internal attention breath-watching meditation compared to the external attention cognitive task in all subjects, indicating a neuro-temporal duality; c) advanced meditators did not show this neuro-temporal duality, as they exhibited no significant difference in the duration of timescales (ACW) between internal (breath-watching) and external (visual oddball) tasks. In other words, they demonstrated what we tentatively describe as ‘*neuro-temporal non-duality*’ (see below for details), while novice meditators and controls did show a significant difference in ACW between these tasks. Lastly, d) non-duality at the psychological level correlated with the difference in ACW between internal and external tasks: a smaller difference in ACW between these tasks was associated with a greater sense of non-duality at the psychological level. Together, our findings highlight the crucial role of the brain’s intrinsic neural timescales in mediating the experience of non-duality in advanced meditators (Figure 6).

**Fig. 6:**
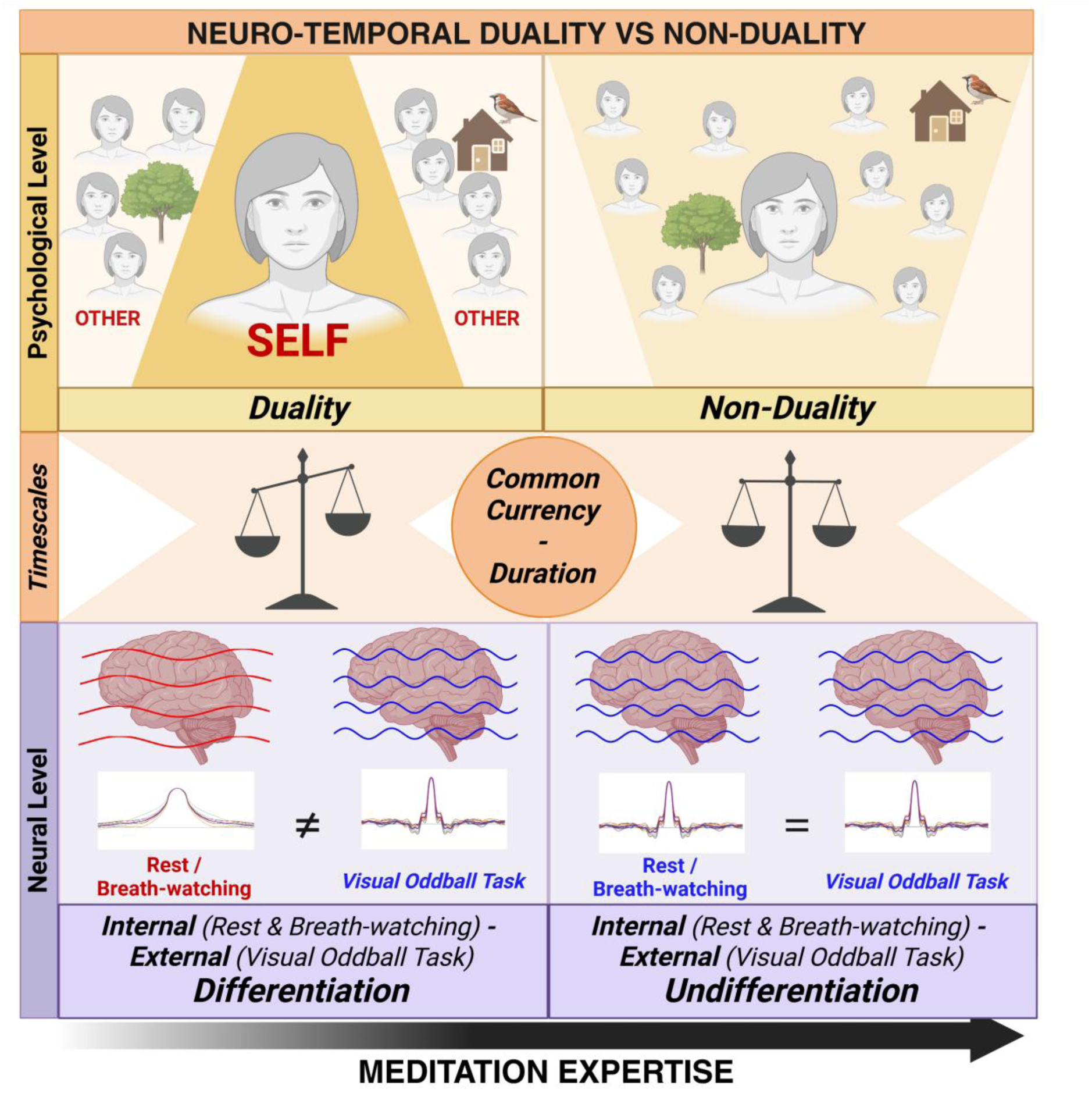
Summary. Our study links the psychological to neural indices of duality and non-duality in study participants. We conceptualize duality in terms of internal and external processing, which we operationalized through distinct attention tasks: breath-watching meditation (internal attention) and a visual oddball cognitive task (external attention). Consequently, we operationalize non-duality as a reduction in the distinction between internal and external attention on both psychological (experience of non-duality) and neural (lack of difference in ACW between internal and external attention). On the neural level, we found internal-external differentiation, or different duration of intrinsic timescales, in novice meditators and controls, whereas we found internal-external undifferentiation, or similar duration of timescales, in advanced meditators. The comprehensive approach we adopted allowed us to connect psychological assessments with neural indicators, demonstrating how non-duality manifests in advanced meditators. These results suggest that duration may be shared by both neural and mental features during non-dual states. Duration is measured at the neural level during internal and external attention—if both show no difference in their neural duration, e.g., ACW, they indicate a non-dual state at the psychological level. Though not measured explicitly here at the psychological level, this suggests that duration may also figure prominently in the experience of non-duality; this is, for instance, supported by subjective reports of shorter durations and timespans in meditative states ^24,95,96^. Though tentative, this suggests that duration may be key in connecting neural and mental states—serving as their “common currency” 1,36,40.

### Non-duality and its psychological features

Our initial research aim was to determine if there were any psychological differences in non-duality among the three groups as measured during breath-watching meditation. We found greater experience of non-duality during breath-watching in advanced meditators compared to other groups. Phenomenology may shed a more detailed light on this finding. Interestingly, breath-watching is a focused internal attention meditation practice that involves maintaining internal attention on the breath while ignoring distractions, such as mind-wandering ^69–73^. In the early stages of this practice, such as in controls, the mind frequently wanders from the breath, requiring significant effort and awareness to refocus. However, this tendency to mind-wander diminishes with increased practice, and attention becomes more focused on the breath ^74–78^. Reduced mind wandering and increased attention focus on the breath have been consistently reported in advanced meditators from different traditions ^69,77–88^. Increased duration of attention on the breath and decreased duration of attention on internal mental contents could thus promote more steady temporal windows of attention in advanced meditators. With increasing expertise in breath-watching, as seen in our advanced meditators, there is a concurrent deidentification from internal mental contents, including one’s thoughts, and a reduction in mind wandering. This process fosters a greater experience of non-dual awareness. Further supporting this, de-identification with one’s thoughts and contents has been shown to be the basis for the experience of non-duality ^6,24,70^.

Remarkably, a growing body of literature shows changes that occur with increasing expertise in meditation ^82,83,89–91^. In particular, it has been reported that advanced practitioners can easily focus on their breath, without significant effort or conscious control. For instance, a functional magnetic resonance imaging (fMRI) study investigating the neural correlates of focused attention meditation in expert Tibetan Buddhist and age-matched novice meditators revealed an interesting pattern where expert meditators engaged lesser attentional networks than novices^92^. This suggests that as expertise increases, the need for conscious control diminishes, supporting the notion that extensive attentional training leads to a state where attention flows effortlessly toward the object of focus, enabling a non-dual state of awareness. This experience of the state of non-duality during breath-watching can be facilitated by the advanced meditator’s increased tendency to non-attach themselves and their mind to any specific contents, that is, increased non-attachment. This is exactly what we observed in our advanced mediators – this is manifest, for instance, in their higher degree of non-dual experience.

### Non-duality and its neural features

After identifying a greater experience of non-duality at the psychological level in advanced meditators, our next question was whether this would also be reflected at the neural level. First, we demonstrated that the psychological duality between internal (breath-watching) and external attention (a cognitive task) also manifests at the neural level. Specifically, the internal attention breath-watching task exhibited longer timescale durations (ACW) than the external attention cognitive task across all subjects.

Intrinsic neural timescales have been shown to provide temporal receptive windows that segregate (temporal segregation) and integrate (temporal integration) inputs of varying durations ^1,39,48–55^. Paying attention to an external stimulus—such as during the cognitive task—is relatively easier and limited to the duration of the external stimulus, which leads to efficient temporal segregation, resulting in a narrower ACW ^37,38,40,61^. In contrast, it is much harder to pay attention internally as the duration of our inner contents (e.g., thoughts, breathing) is longer ^8,57,60,62^. Hence, while focusing on the breath, attention often shifts toward other mental contents, leading to lower temporal segregation and greater temporal integration of the breath with the other mental contents; this leads to a widening of the ACW, as we observed in our data. Similarly so for self-related thoughts which exhibit longer duration (than external contents) and are thus associated with wider ACW than non-self-related thoughts ^57^.

This neuro-temporal duality of the ACW between internal and external tasks was observed in novice meditators and controls. In contrast, advanced meditators did not exhibit such a difference in their ACW, indicating neuro-temporal non-duality. Unlike in the internal attention task (breath-watching), no differences between the groups were observed during the external attention task. This indicates that all participants had similar timescales for external processing as they were all exposed to the same stimuli with the same timing. Accordingly, the absence of an ACW difference between internal and external tasks in advanced meditators stems from a reduction in ACW during the internal breath-watching task. This suggests advanced meditators use shorter neural timescales when practicing breath-watching - this highlights a key role for temporal segregation over temporal integration in focused-attention meditation.

Neural timescales shorten and lengthen with the segregation and integration of stimuli, respectively ^37–39,41^. Focused breath-watching requires the temporal segregation of the breath from distracting stimuli, such as thoughts. This process entails reduced internal attention to mental contents/thoughts and, subsequently, deidentification. In turn, deidentification facilitates the experience of no distinction between internal and external, resulting in non-duality in the advanced meditators. The absence of significant ACW differences between the resting state and breath-watching, as well as between the resting state and the external task in advanced meditators, further supports their experience of a non-dual state, which may already be present during the resting state itself: these subjects no longer make a difference between rest, internal and external states in their ACW which could possibly reflect their deep non-dual processing on the neural level, i.e., neuro-temporal non-duality. Psychologically, that may be manifest in a meditative state-like attitude and composure in the experts even in non-meditative states like the resting state as supported by their higher scores in the non-attachment scale.

### Non-duality and its neuropsychological relationships

The final line of evidence supporting the key role of neural duration for non-duality comes from the significant correlations we observed between the psychological and neural measures of non-duality. Greater experience of non-duality was associated with smaller differences in ACW between the internal attention breath-watching task and the external attention cognitive task. Overall, our findings imply that individuals, through meditative training, can experience various events from a non-dual state of awareness rather than habitually categorizing everything into dualities such as self vs. other, good vs. bad, and internal vs. external. This finding aligns with other studies and reports, indicating a reduction in dualistic distinctions on psychological and neural levels in advanced meditators ^5,6,9–11,18–21,24,26,29,93,94^.

These results suggest that duration may be shared by both neural and mental features during non-dual states. Specifically, duration is measured at the neural level during internal and external attention—if both show no difference in their neural duration, e.g., ACW, they indicate a non-dual state at the psychological level. Though not measured explicitly here at the psychological level, this suggests that duration may also figure prominently in the experience of non-duality; this is, for instance, supported by subjective reports of shorter durations and timespans in meditative states ^24,95,96^. Though tentative, this suggests that duration may be key in connecting neural and mental states—serving as their “common currency” ^1,36,40^.

### Non-duality and its implications

Non-duality, observed cross-culturally, is considered the highest state of being and is part of the phenomenal repertoire of humanity ^12,13,25,70,97–100^. Non-dual awareness has also been referred to as consciousness-as-such, pure consciousness, pure awareness, open awareness, choice-less awareness, turiya, satchitananada, buddha nature, etc.^6,9,12^. Eastern traditions ascribe qualities such as bliss, timelessness, expansiveness, clarity, etc., with experience of non-dual awareness ^9,24,25,70,97^. The experience of non-duality could have significant implications for individual happiness and well-being ^12^. In this context, our findings indicate that non-duality is also part of the neural repertoire of humanity, with meditation serving as a valuable tool for cultivating this state. Our study underscores the value of integrating meditative practices into mental health and well-being programs by demonstrating that advanced meditators show non-duality at both psychological and neural levels. Meditation, a trainable skill, helps to cultivate resilience against everyday stressors and maintain a non-reactive, balanced state of mind ^24,69,72,74,87,101–106^. As the prevalence of mental health issues increases globally ^107–111^, qualities such as non-duality offer a pathway to achieving a deeper sense of peace and stability.

## Limitations

While ACW values suggest a non-dual state in advanced meditators during external tasks, the absence of a psychological measure of duration itself, like duration perception, in this state is a limitation of the current study. Further, future research should incorporate paradigms that elicit stress and emotions to determine whether the non-dual state in advanced meditators is affected by such events. Finally, non-duality is a complex state with several features ^5–8^ which, in addition to the lack of internal-external distinction as investigated here, may warrant future studies. Additionally, one may want to investigate the relationship of non-duality as studied here in relation to some of the traditional mindfulness techniques.

## Conclusion

Our study introduces a novel psychological and neural perspective on non-duality in advanced meditators. First, we show that advanced meditators exhibit greater experience of non-duality during breath-watching meditation. Next, we demonstrate neural differences in auto-correlation window (ACW) duration between internal attention (breath-watching meditation) and external attention (a cognitive task) in novice meditators and controls, indicating neuro-temporal duality. In contrast, advanced meditators no longer show these differences and instead exhibit greater similarity in ACW duration between internal and external tasks, suggesting neuro-temporal non-duality. Strongly supported by correlation findings, we propose that the increased similarity in the brain’s timescales across internal and external tasks serves as a neural marker of non-duality in advanced meditators.

## Acknowledgments

We express thanks to the Isha Foundation volunteers for their support in participant recruitment and sincerely thank Maa Vama, Isha Foundation, for all her support and guidance in conducting this research.

## Methods

### Participants

We carried out additional analyses on our already-published data. The details of all the methods can be found in ref. ^112^. In brief, 103 healthy adults were recruited and categorized into three groups: advanced meditators (ADV), novice meditators (NOV), and meditation-naïve controls (CNT). Advanced meditators (n = 42, 18 females) had a mean (SD) age of 35.57 (6.81) years and a mean (SD) lifetime Isha Yoga practice duration of 5508 (2897) hours. Novice meditators (n = 33, 14 females) had a mean (SD) age of 31.66 (7.64) years with a mean (SD) lifetime practice duration of 1637 (1127) hours in Isha Yoga. Meditation-naïve controls (n = 28, 16 females) had a mean (SD) age of 31.14 (6.38) years and no prior exposure to meditation or yoga schools; they were matched for age, education, and socioeconomic background. This study received approval from the NIMHANS Human Research Ethics Committee (NIMH/DO/ETHICS SUB-COMMITTEE MEETING/2018), and participants provided written informed consent before participation. No monetary compensation was provided.

### Procedure

EEG studies were carried out at NIMHANS. The EEG procedure consisted of four distinct sessions: alternate nostril breathing pranayama (6 minutes), breath-watching (15 minutes), visual oddball cognitive task (15 minutes), and Shoonya meditation (15 minutes) (refer to ref ^112^). Both meditators and meditation-naïve controls underwent these four sessions. Each session included a 4-minute rest period at the beginning and end. Participants were instructed to sit still and relax during these periods. Meditation depth during breath-watching was assessed right after that session.

#### Internal attention (Breath-watching meditation)

Breath-watching is a focused-attention meditation. Controls were given the following instructions: pay attention to the breath, notice it when the mind wanders, and bring it back to the breath. Meditators practiced it as per the Isha Yoga tradition.

#### External attention (visual oddball task)

We utilized a novel paradigm called ANGEL (Assessing Neurocognition via Gamified Experimental Logic) in our study ^113^. This paradigm is based on the visual oddball paradigm, but it includes various gaming elements to increase engagement and allow for the investigation of decision-making in various contexts. ANGEL involves the simultaneous presentation of multiple audio and visual stimuli, challenging participants to make decisions amidst various distractions. Participants received performance feedback at the end of every two blocks to keep them motivated, and instructions and practice sessions were provided prior to the main session.

### EEG acquisition and processing

The data was collected from 128-channel Geodesic EEG System 300 (Philips Neuro, USA). EEG recordings took place in a dimly lit, sound-attenuated chamber with a controlled ambient temperature (25°C) and humidity (40%-60%). Electrode impedance was kept below 50 kohm as per vendor recommendations. Stimulus presentation was managed using E-prime 2.0 software (Psychology Software Tools, Inc., Sharpsburg, PA, USA). Participants were seated 90 cm from a 34 cm × 27 cm LCD monitor displaying instructions. Data were amplified with the NetAmps300 amplifier, digitized at 24-bit resolution, and sampled at 1 kHz (DC amplifier). Electrode positioning followed geodesic spacing guidelines, with Cz as the online reference electrode and AFz as the ground electrode.

The following steps were used in processing the EEG data:

1. Raw EEG data were imported into EEGLAB v2021.0^114^ using the Philips ’mff’ import plugin and pre-processed with in-house MATLAB scripts.
2. The data was resampled to 250 Hz and re-referenced to the average reference.
3. High-pass filtering (>0.5 Hz) and low-pass filtering (<80 Hz) were applied, along with a notch filter at 50 Hz.
4. Bad channels and segments were automatically rejected using the artifact subspace reconstruction (ASR) plugin with a five-standard deviation threshold. On average, 11.19% (min: 1.33%, max: 45.16%) of data were removed for advanced meditators, 12.61% (min: 1.3%, max: 66.7%) for novice meditators, and 11.21% (min: 0.38%, max: 57.17%) for controls.
5. Scalp muscle, ocular, and ECG artifacts were identified and removed using the ICALABEL plugin (90% threshold) after running independent component analysis (ICA) with the ’infomax’ method. On average, 3.14 components (min: 0, max: 21) were removed for advanced meditators, 2.41 components (min: 0, max: 21) for novice meditators, and 3.99 components (min: 0, max: 25) for controls.
6. Channel interpolation was performed using neighbouring electrodes with a spherical spline approach.

### Data cleaning

EEG data were collected from 103 participants. The meditation depth questionnaire (see below) was published ^67^ after data collection had begun and was therefore administered to 70 participants. After data cleaning, 53 subject’s data were available for correlation analyses between EEG measures and the non-duality cluster of the meditation depth questionnaire (see below).

### Autocorrelation window

An autocorrelation function (ACF) was calculated in MATLAB (version 2023b) using a custom code ^115^. This function was applied to 10-second EEG epochs at each electrode with 1-second sliding across relevant session data. Within each 10-second epoch, ACF was computed on 1-second windows (with 50% overlap) and averaged to get a smooth estimate of ACF. On this, ACW-50 was computed as the first lag where the ACF decays to 50% of its maximal value ^39^. Finally, the ACW-50 mean (from 20% trimmed mean) values were calculated from the 10-second epochs for each of the electrodes for each participant. The ACW mean values from the frontal cluster (channels: ’AF8’,’AF4’,’Fp2’,’AFz’,’Fp1’,’AF3’,’AF7’) were considered for the final analysis (Figure 7).

**Fig. 7:**
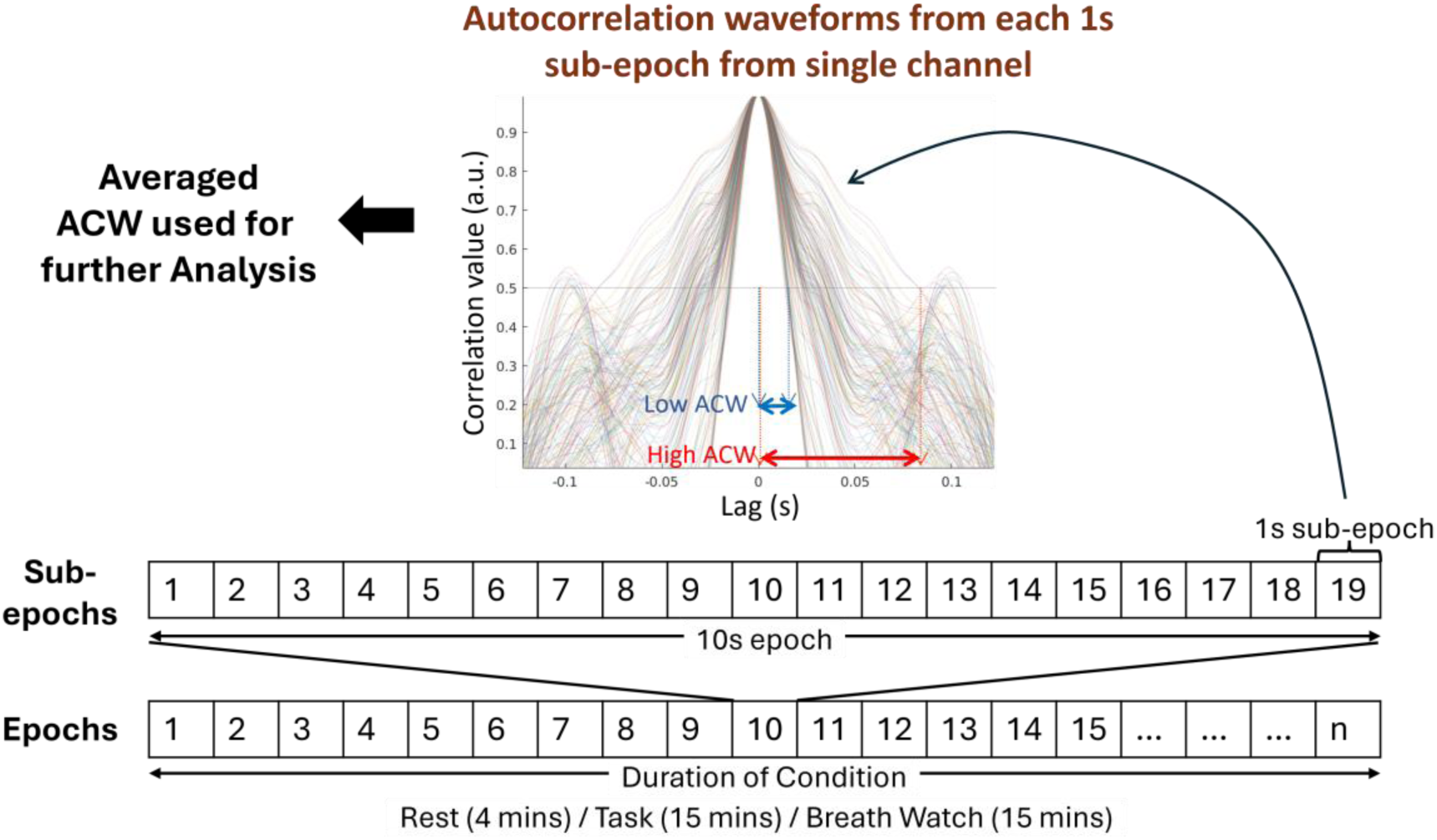
Steps to calculate auto-correlation window (ACW) Steps to calculate auto-correlation function and ACW mean.

### Subjective questionnaires

a. **Non-attachment Scale (NAS-7):** A 7-item version of the original nonattachment scale, known as the NAS-7 ^68^, was utilized to assess nonattachment as a trait feature of the subjects. Participants rated their agreement with seven statements, such as “I can let go of regrets and feelings of dissatisfaction about the past,” using a 7-point Likert scale ranging from 1 (Strongly Disagree) to 7 (Strongly Agree). The internal consistency was good (McDonald’s ω = 0.87, Cronbach’s α = 0.77).
b. **Meditation Depth Questionnaire (MEDEQ):** MEDEQ, a 30-item questionnaire, evaluates the depth of meditation practice ^67^. Participants rate their level of agreement with meditation-related statements on a scale from “0 = not at all” to “4 = very much.” The total score ranges from 0 to 120, with higher scores indicating deeper meditation. The questionnaire comprises five clusters: hindrances, relaxation, concentration, essential qualities, and non-duality. MEDEQ was acquired in 70 participants (controls – 23, novice meditators – 22, advanced meditators – 25). The internal consistency for MEDEQ was excellent (McDonald’s omega = 0.99, Cronbach’s α = 0.96).
c. **The non-duality cluster of MEDEQ:** Items 6 (I felt myself at one with everything), 7 (There was no subject and no object anymore), 16 (Thoughts had come completely to rest), 23 (My mind/consciousness expanded to an infinite space), 25 (There was no differentiation, comparison or judgment anymore. Everything could be as it was) and 26 (My mind, the field of consciousness and awareness was empty from thoughts, emotions and sensations) are part of the well-validated non-duality cluster of the meditation depth questionnaire ^67^. This is a standard instrument for assessing the experience of non-duality, and it has been applied in many studies ^96,112,116–128^. The final score for non-duality is obtained by adding the scores for all these items. A higher score indicates greater non-duality. The internal consistency was good (McDonald’s ω = 0.82, Cronbach’s α = 0.87).

### Data analysis

The statistical analyses were conducted using RStudio Version 1.4.1106, employing several R packages, including ggplot2^129^, ggsci^130^, ggstatsplot^131^, dplyr^132^, effectsize^133^, performance^134^, and car^135^ for necessary analyses. Parametric tests (Student t-test, Mixed ANOVA) or non-parametric tests (Kruskal-Wallis) were used per the normality assumptions. Effect sizes were reported with 95% confidence intervals, and significance was set at p < 0.05. Pearson’s r was used to estimate correlation when both the variables were continuous. Further, Spearman’s rho method was used to estimate the correlation when one of the variables was discrete. Visualization was done using the ggplot library, and each dot was colored as per group. A line of fit was drawn with a 95% CI shadow.

### Replication dataset

We analysed data from 43 expert meditators from our previous study on Vipassana meditation ^136^. The data were acquired with the same EGI 128-channel EEG system employed in the current study. The dataset included senior Vipassana practitioners (n = 22; 11 females; mean age = 54.2 ± 12.6 years; meditation experience = 13.0 ± 4.4 years; cumulative meditation hours = 10,364 ± 5,229) and teachers (n = 21; 10 females; mean age = 51.8 ± 12.2 years; meditation experience = 16.3 ± 5.3 years; cumulative meditation hours = 15,349 ± 9,307). No meditation-naïve control group was included. As part of the experimental protocol, participants performed Anapana (breath-watching meditation) and completed a visual oddball task (ANGEL), among others. Non-duality was not studied. ACW analysis was conducted using the exact same procedure as in the current study.

