## Supplement File for "Non-duality in brain and experience of advanced meditators – Key role for Intrinsic Neural Timescales"

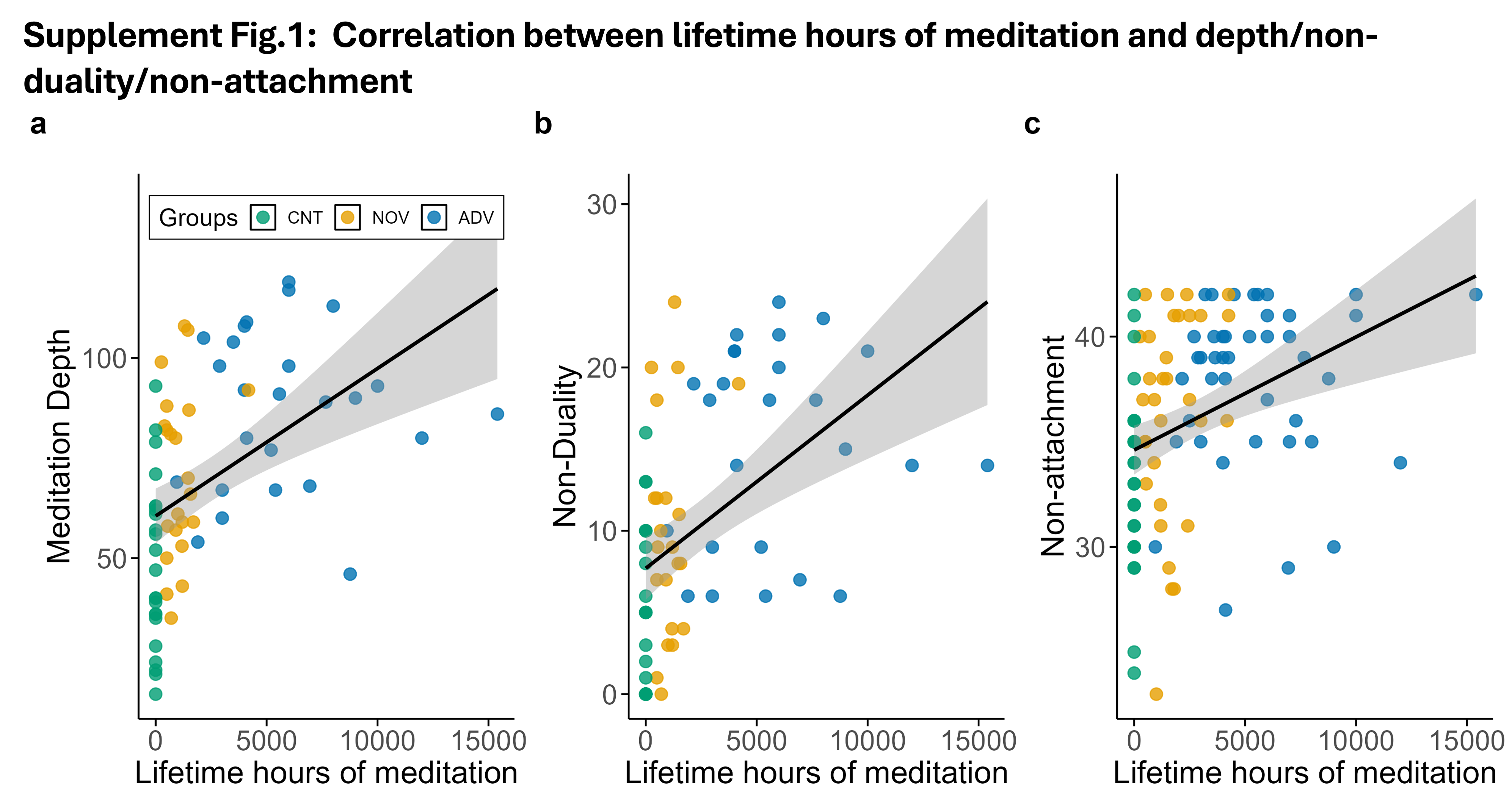

*Scatterplot illustrating the positive correlation between lifetime hours of meditation and* ***a)*** *meditation depth (Ρ_Spearman_* = *0.60*, *p < 0.0001, n = 70),* ***b)*** *non-duality (Ρ_Spearman_* = *0.57*, *p < 0.0001, n = 70), and* ***c)*** *non-attachment (Ρ_Spearman_ = 0.44, p < 0.0001, n = 103).*

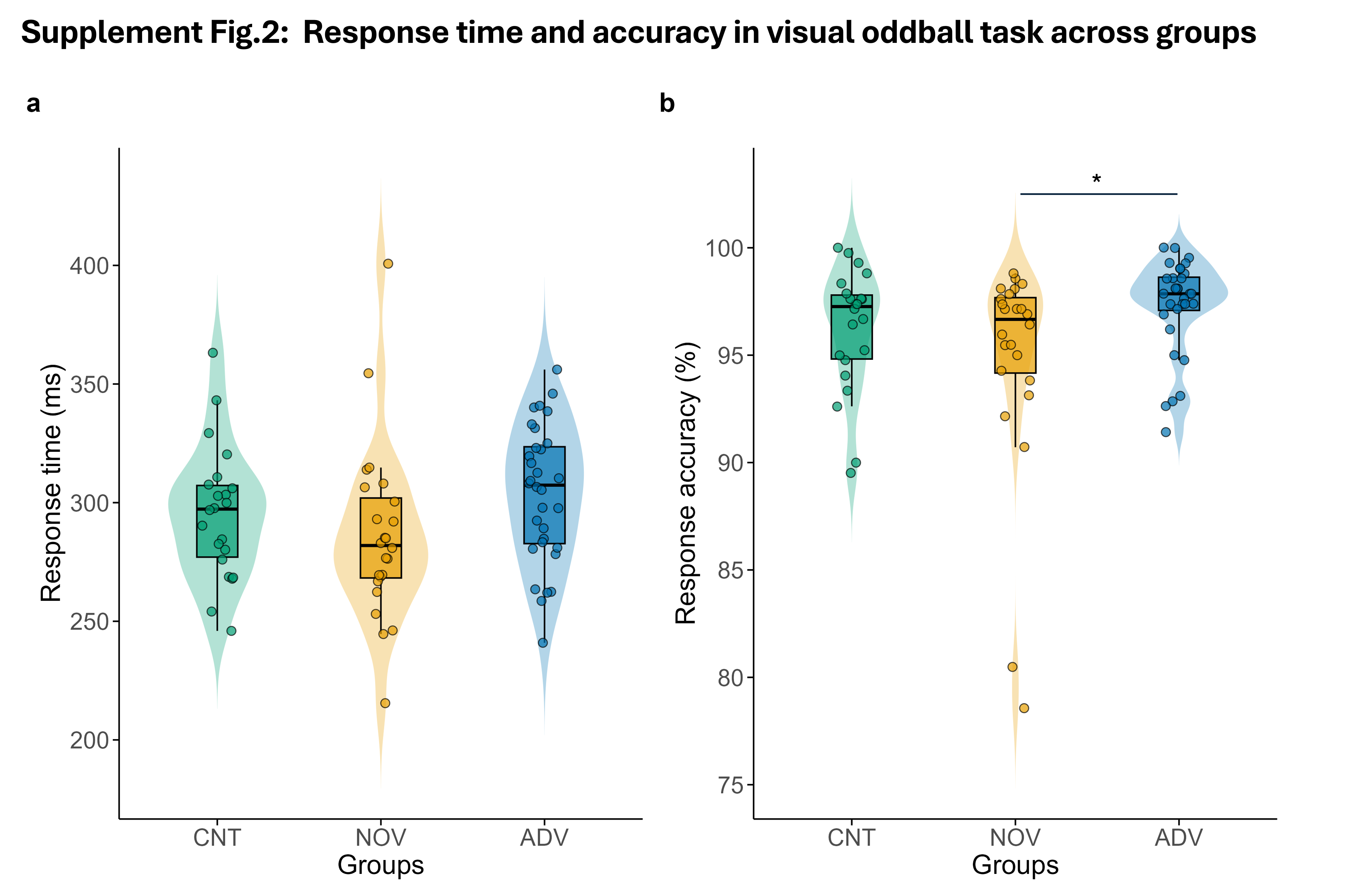

***a)*** *No Significant differences in response time (in milliseconds) across different groups (χ^2^_Kruskal-Wallis_(2) = 5.89, p = 0.05, ϵ^2^_ordinal_ = 0.08, CI_95%_[0.03, 1.00]), n = 78; CNT = 22, NOV = 24; ADV = 32)* ***b)*** *Significant differences in Response accuracy (percentage) across the same groups (χ^2^_Kruskal-Wallis_(2) = 7.36, p = 0.03, ϵ^2^_ordinal_ = 0.10, CI_95%_[0.02, 1.00], n = 78;* *CNT = 22, NOV = 24; ADV = 32). Advanced meditators show highest response accuracy. Significance levels: *p < 0.05. CNT: Controls, NOV: Novice meditators, ADV: Advanced meditators.*

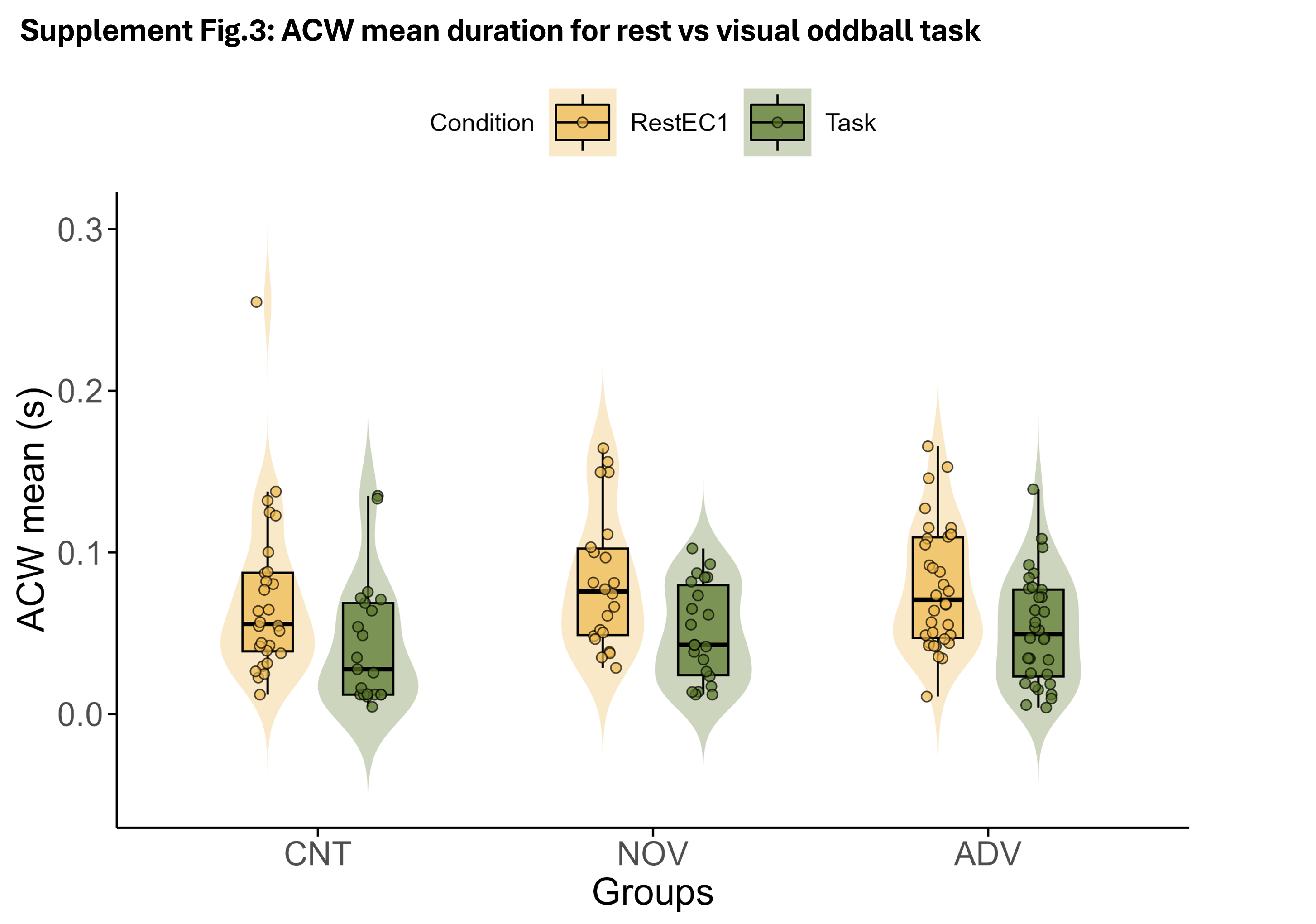

*The difference in ACW mean between eyes closed resting state and visual oddball cognitive task. Statistics are given in Supplementary Table 1.*

**Supplementary Table 1: Results of the Mixed ANOVA Examining the Effects of Group and Condition (Rest, Visual Oddball External Task) on ACW Mean**

| **Effect** | **Sum_Sq** | **df_num** | **Error_SS** | **df_den** | **F_value** | **p_value** |
| --- | --- | --- | --- | --- | --- | --- |
| Intercept | 0.51907 | 1 | 0.125981 | 61 | 251.3328 | < 2.2e-16 **** |
| Groups | 0.00278 | 2 | 0.125981 | 61 | 0.6739 | 0.5135 ^ns^ |
| Condition | 0.02399 | 1 | 0.076166 | 61 | 19.2139 | 0.00004689 **** |
| Interaction (Groups × Condition) | 0.00044 | 2 | 0.076166 | 61 | 0.1753 | 0.8396 ^ns^ |

*This table presents the results of a mixed ANOVA assessing the main effects of Group (Controls, Novice Meditators, Advanced Meditators) and Condition (Rest, Visual oddball cognitive task) and their interaction on ACW mean. The table includes the sum of squares (Sum_Sq), numerator degrees of freedom (df_num), error sum of squares (Error_SS), denominator degrees of freedom (df_den), F-values (F_value), and corresponding p-values (p_value). Significance levels are denoted as: ns = Not significant (p ≥ 0.05), **** = p < 0.0001.*

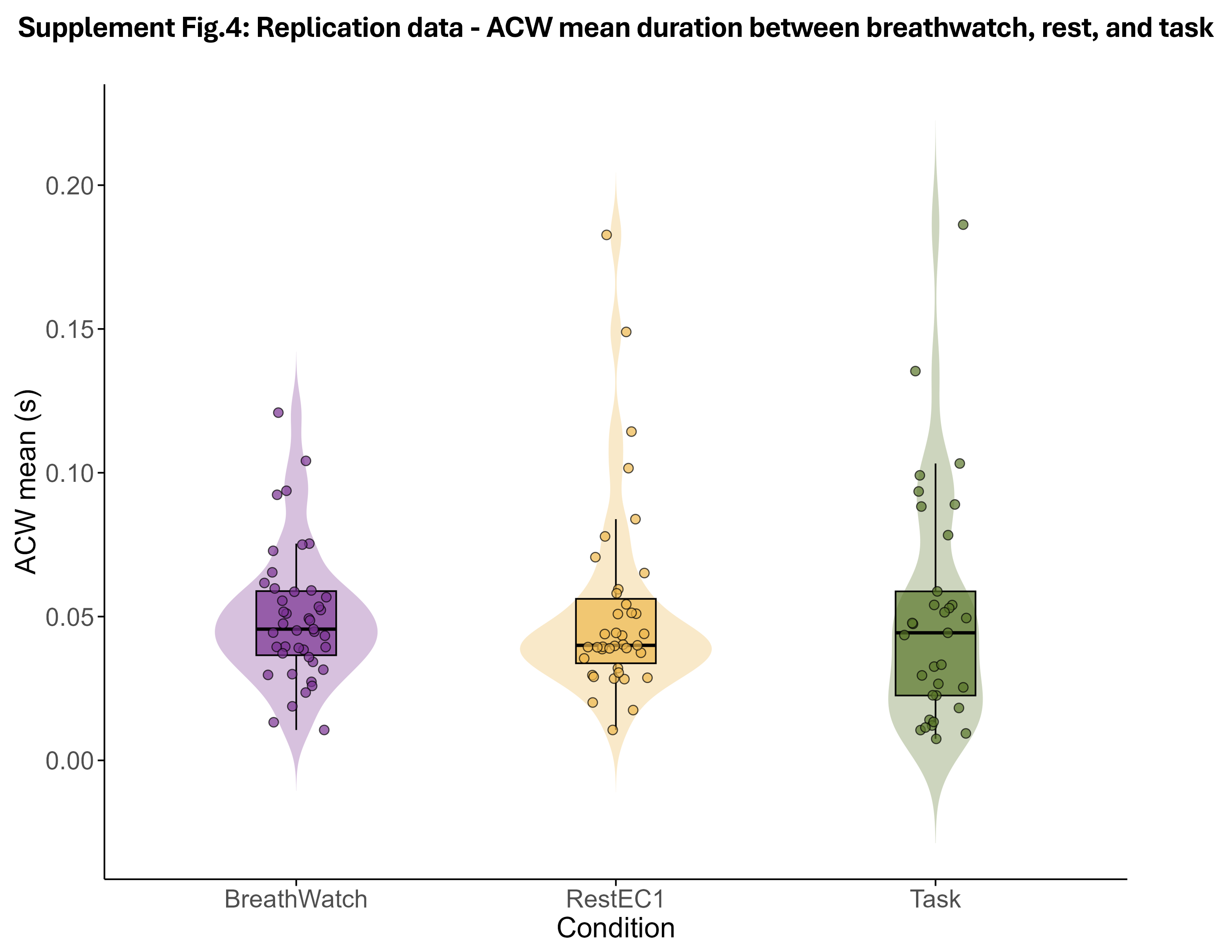

*No significant difference in ACW mean duration between internal-attention breath-watching meditation, rest eyes closed and external-attention visual oddball cognitive task in an independent sample of expert meditators from the Vipassana tradition (F(2, 64.68) = 0.06, p = 0.95, CI_95%_[0.00, 1.00], n = 115; BreathWatch = 43, RestEC1 = 39, Task = 33)). RestEC1 = Rest Eyes Closed.*
